# PARTAGE: Parallel analysis of replication timing and gene expression

**DOI:** 10.1101/2025.09.02.673838

**Authors:** Lakshana Sruthi Sadu Murari, Quinn Dickinson, Silvia Meyer-Nava, Juan Carlos Rivera-Mulia

## Abstract

The human genome is partitioned into functional compartments that replicate at specific times during the S-phase. This temporal program, referred to as replication timing (RT), is co-regulated with the 3D genome organization, is cell type-specific, and changes during development in coordination with gene expression. Moreover, RT alterations are linked to abnormal gene expression, genome instability, and structural variation in multiple diseases, including cancer. However, mechanistic links between RT, large-scale 3D genome architecture, and transcriptional regulation remain poorly understood. A major limitation is that current approaches require the separate profiling of RT and transcriptomes from independent batches of samples, obscuring the complex co-regulation between the epigenome and transcriptome. Here, we developed *PARTAGE*, a multiomics approach that enables joint profiling of copy number variation (CNV), RT, and gene expression from the same sample, providing a more accurate integrative view of the complex relationships between RT and gene regulation.

## INTRODUCTION

Human cells duplicate their genome by the firing of thousands of origins that are activated in clusters following a precise temporal order^1–4^. This replication timing (RT) program partitions the genome into functional units that segregate to distinct nuclear compartments^5,6^. In fact, RT aligns with the 3D genome organization measured by chromosome conformation methods^7,8^, with active and inactive compartments replicating early and late, respectively^5,9,10^. Establishment and maintenance of chromatin epigenetic states also require proper RT regulation^11,12^. RT also changes dynamically during development and is cell type-specific^13–17^. Indeed, around half of the human genome changes RT during differentiation into distinct cell fate specification pathways, and these changes occur in coordination with the establishment of cell type-specific transcriptional programs^15^. Moreover, abnormal RT is linked to altered gene regulation in distinct disease states^18–22^, highlighting the importance of RT control. The correlation between RT and transcriptional activity has been observed for decades, but the mechanistic bases remain elusive^2,5,23,24^. One limitation is the lack of methods enabling the simultaneous measurement of RT and gene expression from the same samples. Although multiple RT profiling methods exist at the ensemble population level^22,25–27^, as well as at the single-cell level^28–32^, links to gene regulation are still being explored by transcriptome analyses on separate sample batches, hindering the complex coregulation of RT and gene expression.

The rapid development of multiomic methods provided tools for the assessment of cell identity and cell-to-cell heterogeneity. Multiomic, single-cell methods also inform on distinct aspects of the epigenome in parallel with gene expression^33–35^. Approaches that map features of 3D genome architecture and transcriptomes have been developed^34,36,37^. However, to date, there is no robust method for the joint mapping of RT and gene expression. In addition, distinct challenges remain to align 3D genome features to gene regulation due to data sparsity, and compartment identification is feasible only with data aggregation^38^. Here, we developed PARTAGE, a multiomic approach for the simultaneous profiling of RT and gene expression. This approach is based on sample preparation under conditions that enable the preservation of both nucleic acids, enabling joint profiling of RT and transcriptome programs. Since this method is performed on asynchronous cultures of cells maintained in regular media conditions, it eliminates the need for cell cycle synchronization, allowing a more accurate measurement of gene regulation. Moreover, the precise collection of cell populations across distinct stages of the cell cycle allows PARTAGE to also provide accurate measurements of CNV. Overall, PARTAGE is a robust approach for the simultaneous analysis of CNV, RT, and gene expression.

## RESULTS

### PARTAGE experimental design

PARTAGE is a multiomics method designed to accurately map CNV, RT, and transcriptome programs from the same samples. It is based on the sample collection for cell cycle analysis under conditions that preserve both nucleic acids. First, cells are labeled with nucleotide analogs (BrdU) to enable the capture of nascent DNA; then, single-cell suspensions are obtained and intact nuclei purified (Figure 1). To enable accurate cell cycle analyses and cell sorting into specific populations while preserving both nucleic acids, intact nuclei are stained with cell-permeant DNA dyes. Specifically, we exploited cell-permeant DyeCycle dyes (Invitrogen), which we previously validated for Repli-seq sample preparation^39^. Then, cell cycle analyses are performed based on DNA content, and fractions of nuclei at distinct stages of the cell cycle are collected by fluorescence-activated cell sorting (FACS). Next, DNA and RNA are co-purified using silica-based strategies and are processed for optimized Repli-seq^22^ and total nuclear RNA-seq. Since PARTAGE enables the isolation of a pure population of nuclei in G1, an accurate CNV mapping can be performed. DNA isolated from S-phase fractions is exploited to measure RT, and the RNA enables accurate detection of transcriptome control during the cell cycle (Figure 1).

**Figure 1.**
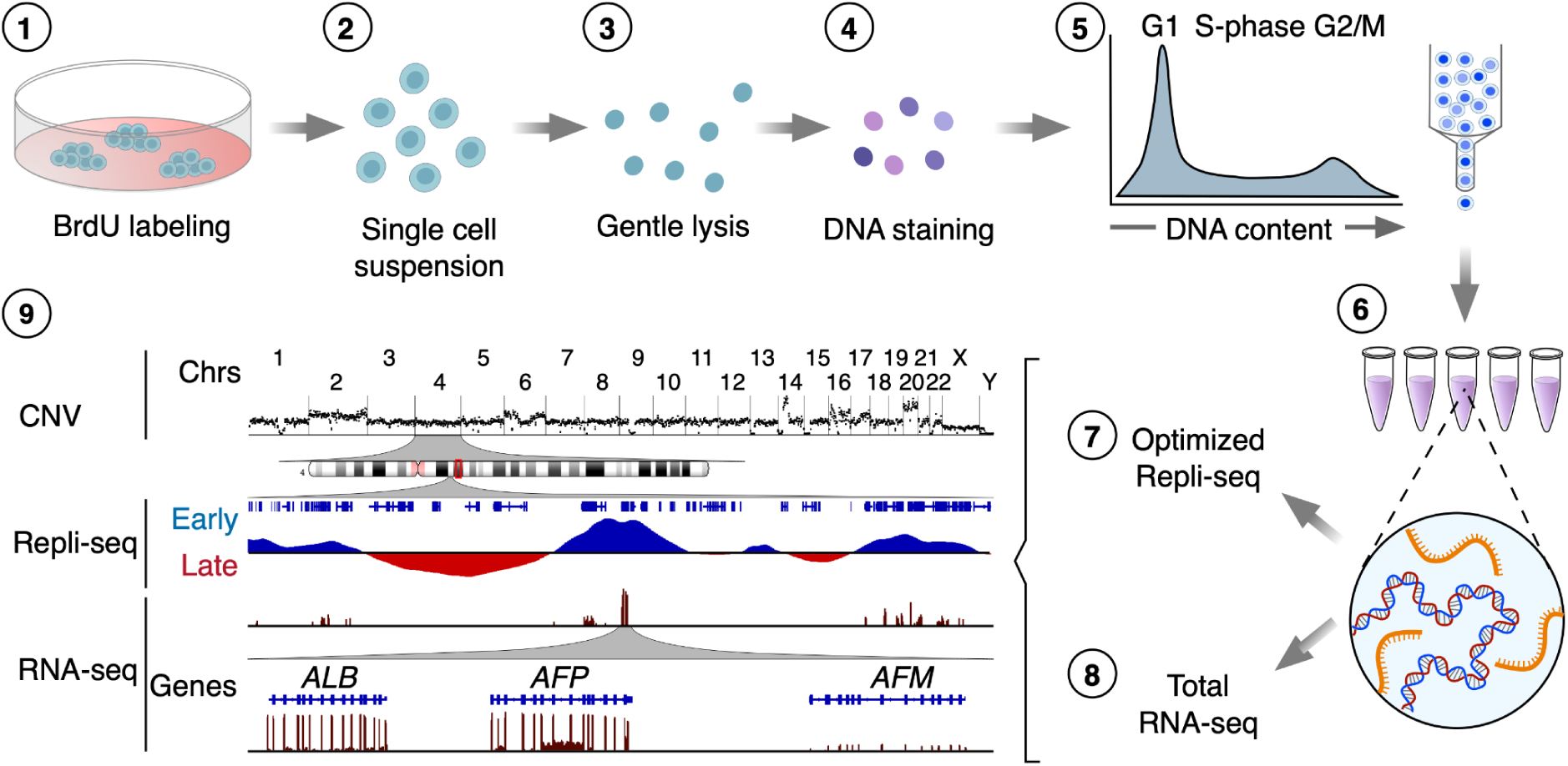
PARTAGE: Parallel analysis of RT and gene expression. Multiomics approach for the simultaneous profiling of CNV, RT, and transcriptome programs. Target cells are BrdU-labeled (1), dissociated (2), and intact nuclei isolated (3). Then, DNA was stained with cell-permeant dyes (4) and cell cycle analyzed by flow cytometry (5). Nuclei at specific cell cycle stages were collected for nucleic acid co-purification (6), and optimized Repli-seq (7) and total RNA-seq (8) were performed. Finally, data analysis and visualization were performed to generate CNV, RT, and gene expression profiles (9).

### PARTAGE sample preparation preserves DNA and RNA integrity

To test PARTAGE, we exploited the HepG2 human cell line, a reference cell line used at the ENCODE^40^ and Roadmap Epigenomics^41^ consortia with extensive omics data publicly available. Cells were grown up to 60% confluence to avoid contact inhibition, which halts normal cell cycle proliferation; then, dissociated into a single-cell suspension, counted, and assessed for viability (Figure 2A, Supplementary Figure 1). Then, intact nuclei were isolated by detergent lysis (Figure 2B, Supplementary Figure 1) and DNA stained for cell cycle analysis (Figure 2C) as previously described^39^. Purified nuclei were analyzed by flow cytometry, and the cell cycle was estimated based on DNA content (Vybrant Violet DyeCycle intensity). Flow cytometry gating strategies and identification of cell cycle fractions are shown in Figure 2D. Populations of 20K nuclei for each fraction derived from G1, S-phase (divided into four fractions), and G2/M were sorted into multiwell plates containing a nucleic acid stabilization solution to preserve integrity. Then, high-quality total genomic DNA and RNA were co-purified using silica-based column methods. PARTAGE sample preparation generated nucleic acid samples of high quality and integrity adequate for downstream processing by optimized Repli-seq and total nuclear RNA-seq (Figure 2E-J, Supplementary Figures 2-3).

**Figure 2.**
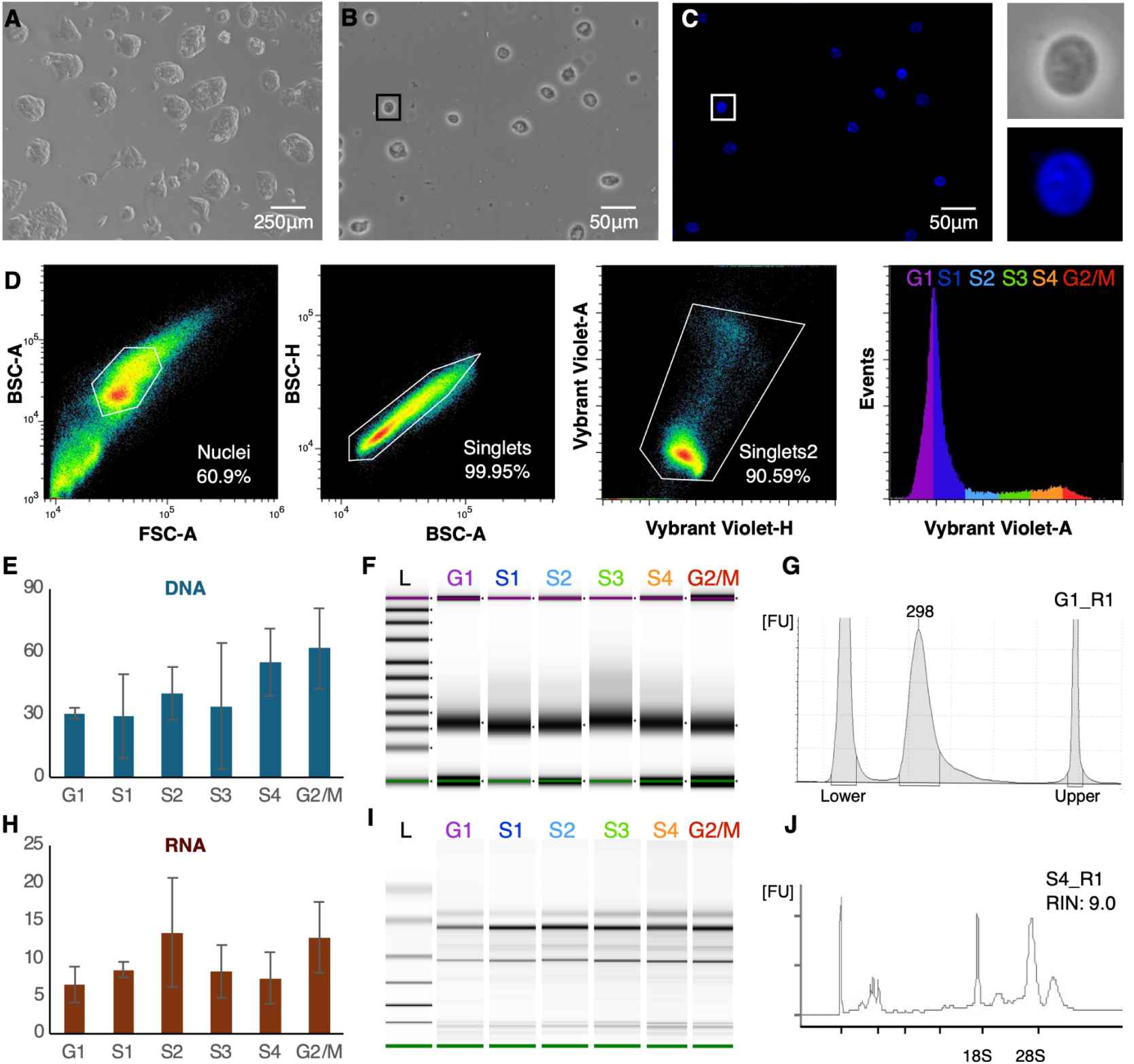
PARTAGE sample preparation. A) HepG2 cells in culture. B) Purified intact nuclei. C) Nuclei staining with Vybrant DyeCycle Violet for accurate DNA content estimation. Magnification of a single nucleus is shown. D) Flow cytometry and gating strategy. Single nuclei were identified and gates were established to purify cell populations across the cell cycle: G1, S1, S2, S3, S4, G2/M. 20K nuclei were collected per fraction, and three independent biological replicates were processed. E) DNA yields (total ng) obtained from each sorted population. F) Integrity analysis and size distribution of purified and fragmented genomic DNA. G) Electropherogram of purified and fragmented genomic DNA. H) RNA yields co-purified with the DNA (total ng) from the sorted cell populations. I) RNA integrity analysis. J) Exemplary electropherogram of purified RNA demonstrating RNA integrity (RIN = 0.90).

### PARTAGE simultaneously maps CNV, RT, and transcriptomes

To evaluate PARTAGE’s capacity to capture the multimodal genomic signals from the same samples, we analyzed HepG2 cells and compared the results against those generated using standard Repli-seq and RNA-seq methods (Figure 3). Standard Repli-seq was performed on HepG2 nuclei as previously described^22^. For standard RNA-seq, RNA was isolated from 0.5 million cells, and libraries were prepared for stranded mRNA sequencing. These standard analyses confirmed the close alignment of RT programs with genome compartmentalization as measured by HiC, as well as enrichment of active gene expression in active-early replicating genomic compartments (Figure 3A). However, these datasets were generated independently from separate batches of samples processed by the standard methods of Repli-seq, HiC, and RNA-seq (Figure 3A). Thus, we performed PARTAGE by processing the purified genomic DNA using the optimized Repli-seq method and the RNA using an ultra-low input library preparation pipeline. PARTAGE allowed us to simultaneously map CNV, RT, and gene expression from the same sample (Figure 3B). First, whole-genome sequencing of the cell populations isolated from G1 enabled the accurate mapping of CNV and the detection of multiple amplifications in the HepG2 genome (Figure 3B). Next, RT was mapped using the multifraction Repli-seq method^42,43^. Standard Repli-seq involves the sorting of early and late fractions of S-phase cells/nuclei, and the generation of binary profiles of early versus late replicating values across the genome. In contrast, multifraction Repli-seq involves cell sorting of multiple fractions across the entire S-phase to increase resolution and fine-scale detection of genomic features of the replication domains^43^. Thus, PARTAGE multifraction Repli-seq was performed and data visualized as heatmaps of normalized sequencing signals per genomic bin. We found that the patterns of early/late replication reproduced the RT timing data obtained with standard Repli-seq (Figure 3B). Finally, PARTAGE total-nuclear RNA-seq reproduced the transcriptome of the HepG2 cells obtained from separate RNA-seq analysis (Figure 3). To display global trends in gene expression, we displayed exemplary patterns in three distinct chromosomal regions. To confirm cross-protocol consistency in transcriptomes, we displayed zoomed-in tracks focused on hepatic-specific and housekeeping genes (Figure 3B).

**Figure 3.**
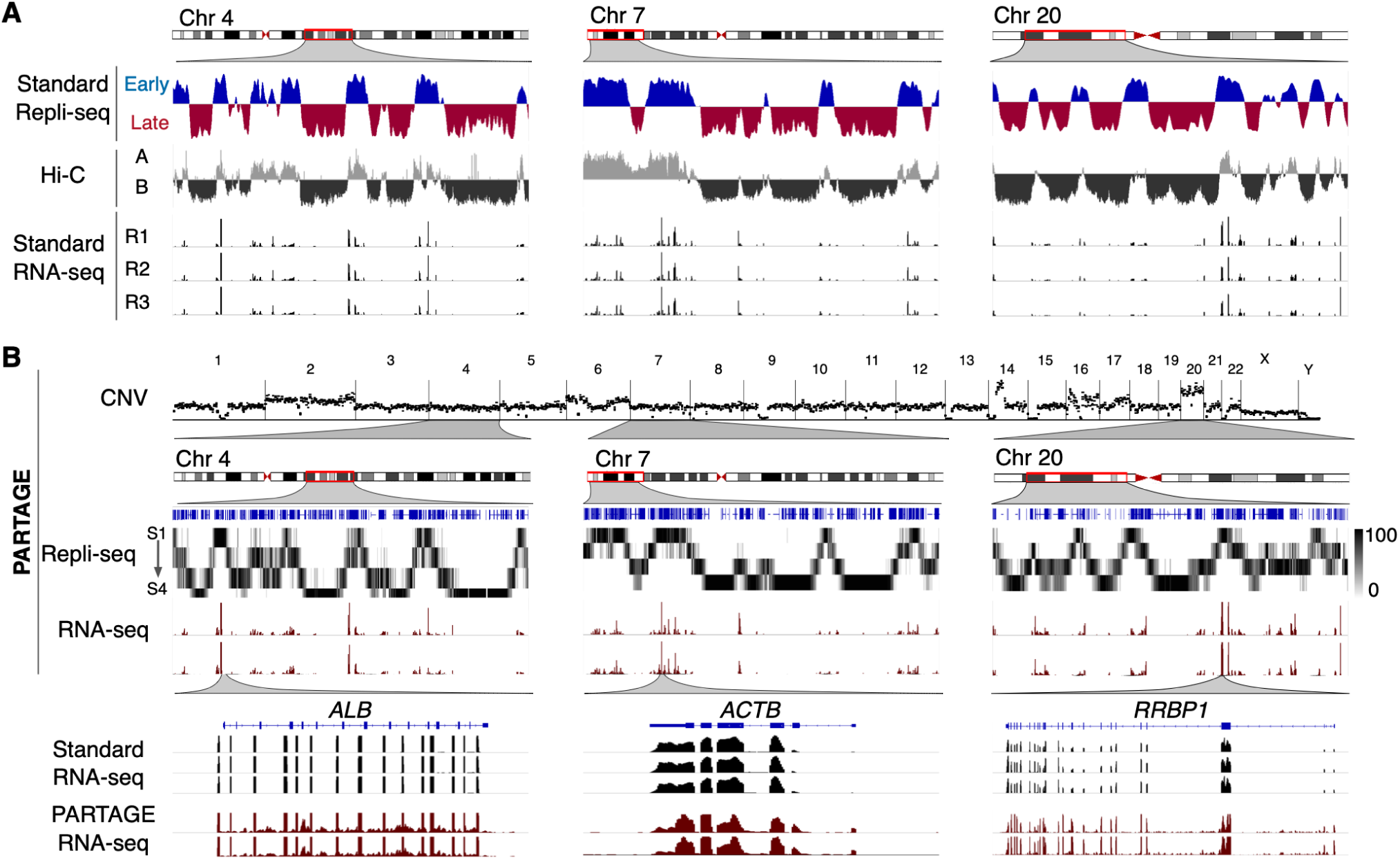
PARTAGE captures CNV, RT, and transcriptomes from the same samples. A) Standard omics analysis generated from HepG2 cells. Three exemplary chromosomal regions are shown. Standard Repli-seq, chromosome confirmation capture (HiC), and RNA-seq were performed from separate sample batches of HepG2 cells. B) PARTAGE simultaneous mapping of CNV, RT, and transcriptome from HepG2 nuclei. CNV was calculated from the sequencing reads obtained from G1 cell populations. Sequencing reads normalized for depth and per 1 Mb bin are shown. PARTAGE Repli-seq was performed using multifraction RT analysis. Heatmap signals of G1-normalized reads are shown per fraction (S1-S4). RNA-seq coverage tracks are shown for the same chromosomal regions. Bottom tracks show a comparison between standard and PARTAGE RNA-seq signals zoomed in on hepatic-specific (*ALB*) and housekeeping genes (*ACTB* and *RRBP1*).

To validate reproducibility and accuracy, we performed in-depth comparisons of PARTAGE multiomic signals against those generated by the separate standard methods. First, the HepG2 genome contains well-known amplifications, and public CNV data exists; thus, we compared PARTAGE-CNV data to the ENCODE CNV data^44^. We found that all major genomic amplifications were identified and aligned with the reference data (Figure 4A). Second, we compared PARTAGE RT programs versus standard Repli-seq and found that the multifraction PARTAGE Repli-seq signals faithfully recapitulated the patterns of the standard Repli-seq (Figure 4B). To enable direct comparison of these different Repli-seq methods, we collapsed the PARTAGE multifraction Repli-seq into log2(Early/Late) ratios. The overlay of RT profiles demonstrated the consistency between PARTAGE and standard Repli-seq (Figure 4B). Moreover, cross-protocol scatterplot comparisons confirmed close concordance of RT values across the genome (Figure 4C). Furthermore, pair-wise/genome-wide correlation analysis confirmed high consistency between PARTAGE and standard Repli-seq RT profiles with *r* ≥ 0.95 (Figure 4C-D). Finally, we also validated the concordance between PARTAGE and standard RNA-seq pipelines to ensure accurate transcriptome profiling was achieved in this method. RNA-seq raw coverage tracks (normalized to sequencing depth) demonstrated that PARTAGE recapitulates the transcriptome data obtained from standard methods at specific genes (Figures 3B and 4E). Since transcriptomes were obtained from distinct cell numbers (0.5 million cells for standard RNA-seq versus 20,000 nuclei in PARTAGE RNA-seq), and using distinct library preparation pipelines (TruSeq stranded RNA-seq versus Takara ultra-low input SMARTer Stranded Total RNA-Seq), we scaled and normalized the data for proper comparison (Supplementary Figure 4). Then, Log2TPM RNA-seq values were median-centered and batch corrected^45^ to enable proper comparisons. We found strong gene-wise concordance between PARTAGE and standard RNA-seq, and replicate-level scatterplots of gene expression values showed near-isometric agreement, with housekeeping and hepatic-specific marker genes aligning closely (Figure 4F, Supplementary Figure 5). Moreover, Pearson’s correlation across all standard/PARTAGE replicate pairs resulted in *r* ≥ 0.945 (Figure 4G), indicating strong reproducibility between the distinct RNA-seq library preparation pipelines and accurate transcriptome profiling in PARTAGE. Overall, these results validate the simultaneous and precise mapping of CNV, RT, and gene expression patterns using PARTAGE.

**Figure 4.**
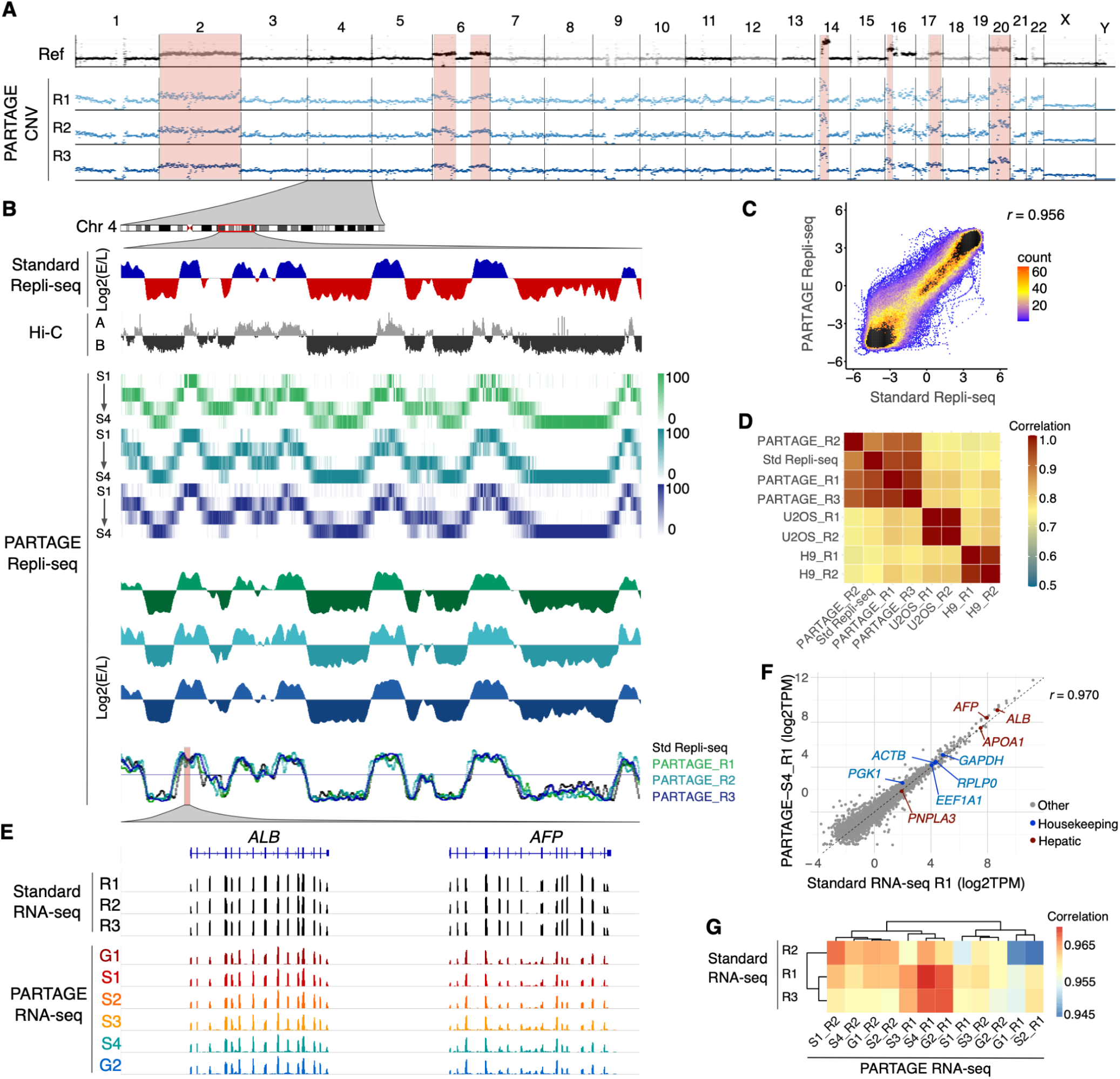
PARTAGE accurately profiles CNV, RT, and transcriptomes. A) CNV analysis using cell samples collected from G1. Reference track (ref) was obtained from the ENCODE data portal. Amplified chromosomal regions in the HepG2 genome are highlighted. B) Comparative analysis of standard and PARTAGE Repli-seq data. Standard Repli-seq is shown as log2(E/L) ratio signals. Multifraction Repli-seq is shown in distinct heatmap colors per replicate. Signals are normalized counts against G1 per 20kb bins. Collapsed E/L Log2 ratios were calculated from PARTAGE multifraction Repli-seq RT to obtain RT profiles directly comparable to standard E/L Repli-seq. C) Scatterplot of PARTAGE (y-axis) versus standard (x-axis) Repli-seq data. D) Genome-wide correlation between a PARTAGE and standard Repli-seq. Genome-wide RT data in 20 Kb bins were used. RT from stem cells (H9) and cancer cells (U2OS) were included as a comparison. E) PARTAGE and standard RNA-seq coverage tracks at the albumin locus showing expression patterns of the hepatic genes *ALB* and *AFP*. Raw coverage normalized for sequencing depth was used. F) Gene-by-gene scatterplots on batch-corrected expression values (log2TPM). Standard RNA-seq (x-axis) versus PARTAGE RNA-seq (y-axis) is shown. Housekeeping genes (blue) and hepatic markers (red) are highlighted; all other genes are gray. G) Genome-wide cross-protocol concordance between a PARTAGE and standard RNA-seq transcriptomes. Pearson correlation among all samples, computed on batch-corrected Log2TPM values, and clustered by Euclidean distance, is shown.

### Cell cycle dynamics in RT and gene expression

A long-standing correlation has been observed between early replication and active gene expression^2,5,14–16^. However, previous studies were based on RT and gene expression data obtained independently from distinct batches of samples generated in parallel. Thus, we exploited PARTAGE to explore how closely early replication and gene expression are linked, taking advantage of the RT and transcriptome measurements from the same sample that reduces technical noise and batch effects. Genomic tracks of PARTAGE RT profiles and transcriptomic coverage patterns confirmed that active transcription is enriched at early replication chromosomal regions (Figure 5A). To measure the strength of this correlation, we calculated the percentage of expressed genes in relation to their RT. To do so, we identified all expressed genes and plotted them according to their RT (collapsed Log2E/L ratios). Our results confirmed a strong enrichment of expressed genes in early replicating regions (Figure 5B). Moreover, a regression analysis was performed and showed a strong correlation between RT and transcriptional activity (Figure 5B). These results confirm the co-regulation between RT and active transcription.

**Figure 5.**
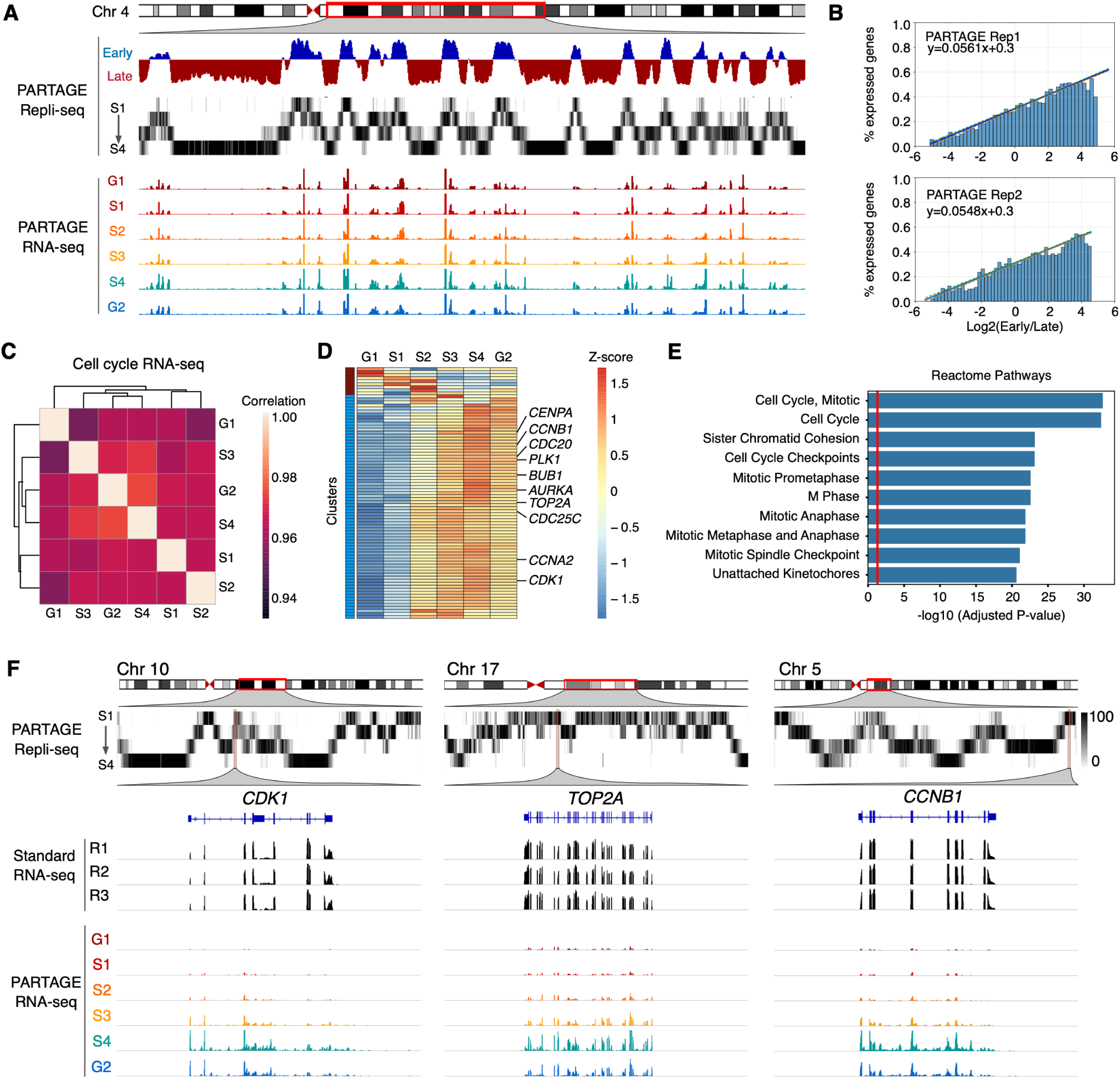
Cell cycle regulation of DNA replication and gene expression. A) Exemplary genomic locus from chromosome 4 showing enrichment of gene expression at the early replicating regions. Collapsed E/L ratios from PARTAGE Repli-seq data, PARTAGE multifraction signals, and coverage RNA-seq tracks are shown. B) Genome-wide correlation between replication timing and gene expression. Expressed genes were identified (threshold of ≥2 log2TPM) and sorted by their RT. The y-axis represents the percentage of expressed genes within each bin. Logistic regression (inner line) and 95% confidence intervals (outer lines) display the correlation strength. Two PARTAGE replicates are shown. C) Genome-wide correlation of transcriptome programs across the cell cycle. Average TPMs were calculated per cell cycle stage after batch correction (ComBat), and gene expression values ≥ 0.5 log2TPM in at least three cell cycle stages were used. Pair-wise Pearson’s correlation was performed with hierarchical clustering based on correlation distance (1-*r*). D) Heatmap and clustering analysis of variable genes during the cell cycle. Genes are represented by row-wise z-scored log₂(TPM) values and clustered by k-means (k = 2). Only genes expressed in ≥2 cell cycle stages and with ≥ 4-fold Δlog2TPM were considered. E) Reactome ontology analysis of the major cluster (blue) is shown. F) Exemplary chromosome regions harboring variable genes that are induced during cell cycle progression (*CDK1*, *TOP2A*, and *CCNB1*).

PARTAGE also enables the profiling of cell cycle phase-specific transcriptomes, enabling direct measurement of dynamic regulation of gene expression patterns. Thus, we analyzed whether changes in gene regulation occurred as the cells progressed through the cell cycle. First, we determined the degree of similarity between the transcriptomes obtained from the distinct stages of the cell cycle. We removed non-expressed genes, applied batch correction, and performed correlation analysis. We found strong correlation coefficients (*r* ≥ 0.95) between all pair-wise comparisons of cell cycle stages (Figure 5C), demonstrating the robustness of PARTAGE and global stability of the transcriptome of HepG2 across the cell cycle. To identify transcriptional changes during the cell cycle, we identified the expressed genes in a minimum of 2 samples and filtered those with ≥ 4-fold change across cell cycle stages. These stringent criteria allow us to identify the robust transcriptional changes during the cell cycle, and we detected 83 protein-coding genes. Clustering analysis revealed a group of co-regulated genes that increase expression during cell cycle progression, containing cell cycle regulators (Figure 5D). Moreover, ontology analysis confirmed that variant genes are associated with processes regulating the cell cycle progression (Figure 5E). Exemplary PARTAGE tracks of cell cycle-regulated genes are shown in Figure 5F. Overall, PARTAGE provides an integrated framework with the potential to detect CNV, RT, and gene expression regulation during the cell cycle.

## DISCUSSION

Analyses of genome organization and function have unveiled the co-regulation between RT and gene expression^15,16,46,47^. Moreover, recent studies have shed light on the RT emergence during the earliest developmental stages and suggest that RT precedes the establishment of the chromatin states and transcriptomes^48–51^. However, all previous studies have been performed by the separate profiling of these genomic properties, which might obscure the complex coregulation between the epigenome and transcriptome. In this work, we present PARTAGE, a multiomics approach that enables the parallel analysis of RT and gene expression, as well as CNV, from the same samples.

The major limitation for the joint mapping of transcriptomes and RT was that methods for cell cycle analysis by FACS relied on methods that do not preserve the RNA. Standard cell cycle analysis by flow cytometry depends on the precise DNA content measurement using propidium iodide (PI) or diamidino-2-phenylindole (DAPI) staining. PI staining requires RNase treatment, while DAPI staining requires cell permeabilization, which leads to RNA loss. To overcome this challenge in PARTAGE, we exploited non-toxic, cell-permeant dyes with narrow emission spectra optimized for flow cytometry (DyeCycle dyes, Invitrogen). These dyes enable the stoichiometric DNA staining for accurate cell cycle analysis by FACS in live cells^52^,, and we confirmed that they can be used for Repli-seq sample preparation^39^. Since these dyes do not need permeabilization and are not RNA-sensitive, they can be exploited for accurate collection of cells from specific stages of the cell cycle by flow cytometry while preserving both nucleic acids. Moreover, whole live cells as well as intact nuclei processing are compatible with this approach. Here, we processed intact nuclei to better capture transcriptional activity. However, PARTAGE can be easily adapted for whole cells for analysis of steady-state RNA-seq. We demonstrated that PARTAGE sample preparation allows the co-purification of DNA and RNA of high quality and in sufficient amounts for parallel processing for Repli-seq and RNA-seq. In fact, we found increasing yields of DNA that reach 2-fold in G2/M as compared to G1, highlighting the accuracy of this method to collect cell populations throughout the cell cycle (Figure 2).

Conventional methods for CNV detection rely on deep sequencing (30X coverage) to accurately identify variations by ensemble methods. Here, we demonstrated that PARTAGE can accurately detect major amplifications in the HepG2 genome with only ≥0.5X coverage. Despite the shallow sequencing (∼8M single-end 100bp reads), PARTAGE reproduced the large CNV detected by the ENCODE mapping (Figure 4). This was achieved by the focused analysis of purified nuclei in G1-phase, which ensures the analysis of uniform diploid populations by removing S-phase and G2/M-phase nuclei, and reducing cell cycle-associated artifacts in CNV detection. Thus, PARTAGE enables the accurate detection of CNV, providing additional tools to better understand genome instability in development, disease, and cancer states. Importantly, PARTAGE’s capacity to detect CNVs with this sequencing depth makes this approach cost-effective and scalable for clonal tracking, genome integrity screenings, and cancer alterations studies, with the addition of joint profiling of RT and gene regulation.

For Repli-seq analysis in PARTAGE, we took advantage of our optimized Repli-seq workflow, which reduced DNA loss and technical noise by minimizing transfer/handling steps, and increasing yields by 3-fold ^22^. We also applied the multifraction Repli-seq approach and exploited the data from the G1-purified nuclei to normalize signals and generate accurate heatmaps of RT. Although standard E/L Repli-seq is also compatible with our PARTAGE workflow, it is sufficient to accurately detect RT, and is more cost-effective; we selected multifraction Repli-seq to enable a more accurate detection of cell cycle regulation when integrating the data with the matching transcriptomes. The combined implementation of multifraction analyses and optimized library preparation strategies allowed us to generate highly reproducible RT data across replicates that recapitulate the RT data obtained from the standard method.

We also evaluated the robustness of the RNA-seq data generated in PARTAGE and found high reproducibility across replicates. Moreover, although PARTAGE RNA-seq was performed on low-input samples (20K nuclei), it faithfully recapitulated the transcriptome obtained with standard RNA-seq from 0.5M cells (Figure 4). Moreover, PARTAGE sample preparation allows us to collect nuclei at distinct phases of the cell cycle. Thus, accurately mapping the cell-cycle regulation of the transcriptome program can be performed. Notably, PARTAGE profiles RNA in native conditions without the need of forced cell synchronization. Conventional studies of cell cycle-regulated transcriptional activity depend on synchronization strategies through serum starvation, mitotic shake-off, or using drugs that activate checkpoint mechanisms. However, these approaches trigger cellular stress responses, DNA damage pathways, and/or metabolic reprogramming that alter normal transcriptional regulation^53–55^. Thus, PARTAGE is a valuable alternative that does not require any external perturbation and enables a more accurate assessment of the cell cycle regulation of the transcriptome. Indeed, the simultaneous profiling of RT and transcriptomes in PARTAGE facilitated the integration of multimodal signals under physiological conditions and confirmed the strong enrichment of actively transcribed genes in early replicating regions (Figure 5). Moreover, PARTAGE identified a subset of genes that increase transcriptional levels 4-fold as the cells progress in the cell cycle, and confirmed that these genes are indeed involved in cell cycle regulation. The relatively small number of cell cycle-regulated genes detected in PARTAGE can be due to different factors: 1) the reduced synchronization-induced artifacts by processing cells in native conditions; 2) the processing of nuclei instead of whole cells to remove bias from post-transcriptional regulation; and 4) the use of stringent thresholds to identify variable genes.

Overall, PARTAGE is a highly robust method that provides accurate mapping of both RT and transcriptomes. This method reduces the required sample input as compared to the corresponding separate omic approaches, reduces heterogeneity and technical noise between distinct batches of samples, and enables a direct assessment and integration of multimodal signals.

## METHODS

### Cell culture

The HepG2 cell line was obtained from ATCC and expanded according to the supplier’s protocol. Briefly, cells were maintained in Eagle’s Minimum Essential Medium (ATCC Catalog No. 30-2003) supplemented with 10% FBS (Corning Cat. No.35011CV). Cell cultures up to 60% confluency were treated with BrdU (Sigma, cat. no. B5002) at a final concentration of 100 µM at 37 °C for 2 hours and processed for nuclei preparation for PARTAGE analyses.

### Intact nuclei preparation for PARTAGE

BrdU-labeled HepG2 cells were dissociated into single cell suspensions by treatment with trypsin-EDTA (Life Technologies, cat. no. A011105-01), and intact nuclei were isolated as previously described^39^. Briefly, cell pellets were resuspended in 1 ml of DMEM/F-12 50/50 (1X), counted, and aliquoted into batches of 2.0 × 10^6^ cells. Then cells were triturated in a lysis buffer: 0.025% IGEPAL (Millipore Sigma cat. no. 542334) in 10mM TrisHCl, 10mM NaCl, 3mM MgCl2, and 1X PBS (Thermo Fisher, cat. no. 20012027). Lysed cells were washed three times and resuspended in 1 mL PBS/1% FBS. Finally, nuclei were stained with Vybrant DyeCycle Violet Stain (ThermoFisher, Cat. #V35003) at 1.5 µl per 1 × 10^6 cells for 30 minutes at 37°C and filtered with a 36-µm nylon mesh (Fisher Scientific, cat no. NC05156770).

### Cell cycle analysis and cell sorting by flow cytometry

Stained nuclei were processed in a SONY SH800 cell sorter. Single nuclei were identified and cell cycle stages determined based on DNA content (Vybrant DyeCycle Violet intensity). 20K cells were collected per fraction of G1, S1, S2, S3, S4, and G2M cell cycle stages and sorted directly into wells of a 96-well plate containing (Axygen, Cat. #14-223-345) 400 μL of DNA/RNA Shield Buffer (Zymo, cat. no. R1100). If not processed immediately, the plates were sealed with an adhesive film and stored at -80 °C.

### Nucleic acid extraction

For DNA and RNA co-purification, the samples were thawed at room temperature and nucleic acids isolated using the Quick DNA/RNA MicroPrep Plus Kit (Zymo, Cat. #D7005) according to the manufacturer’s instructions. Purified DNA and RNA were quantified using Qubit kits (Invitrogen, Cat. Nos. Q32852 and Q32851).

### Repli-Seq

DNA samples were processed using the optimized Repli-seq protocol as previously described^22^. Briefly, purified DNA from sorted cells was directly processed using the KAPA HyperPlus library preparation kit (Roche Cat. No. 07958978001) according to the manufacturer’s instructions. Enzymatic fragmentation was performed at 37 °C for 30 min, followed by end-repair and ligation of KAPA dual-indexed adapters (Roche Cat. No. 08278555702). Libraries were purified using the Zymo DNA Clean & Concentrator-5 kit and processed for BrdU-IP. Immunoprecipitated libraries were amplified using the KAPA HiFi HotStart from the KAPA HyperPlus kit according to the manufacturer’s instructions and purified using AMPure XP beads. DNA concentration and size distribution were evaluated by Qubit quantification (Invitrogen, Cat. No. Q32851), and size estimation was performed on TapeStation High Sensitivity DNA ScreenTape (Agilent 5067-5592). Repli-seq libraries were pooled at a final concentration of 10nM and sequenced at the University of Minnesota Genomics Center (UMGC), on the Illumina NextSeq P2 Illumina platform at ∼8 M reads per library and 100 bp of single-end sequencing.

### Standard RNA-Seq

RNA samples were isolated from 0.5 million HepG2 cells using the RNeasy Plus Kit (Qiagen 74004), and libraries were prepared with the TruSeq Stranded mRNA Library Prep (Illumina) according to the manufacturer’s instructions. Standard RNA-seq libraries were sequenced on the Illumina NovaSeq platform at ∼20 M reads per library and 150 bp of pair-end sequencing.

### PARTAGE RNA-Seq

RNA samples were processed using the SMARTer Stranded Total RNA-Seq Pico for ultra-low inputs (Takara Cat. No. 634411). RNA concentration and size distribution were evaluated by Qubit quantification (Invitrogen, Cat. No. Q32852) and size measured using the Bioanalyzer RNA 6000 Nano Kit (Agilent Cat. No. 5067-1511). RNA-seq libraries were pooled at a final concentration of 10nM and sequenced at the University of Minnesota Genomics Center (UMGC), on the Illumina NovaSeqX platform at ∼40 M reads per library and 150 bp of pair-end sequencing.

### Data analysis and visualization

Data analysis was performed with resources at the Minnesota Supercomputing Institute (MSI).

For PARTAGE CNV data, sequencing reads obtained from G1 fractions were trimmed for adapter sequenced using Cutadapt^56^ and aligned to the reference genome with Burrows-Wheeler Aligner^57^. Sequence Alignment Map (SAM) files were converted into Binary Alignment Maps (BAM) with SAMtools^58^. Quality filter and removal of PCR duplicates were performed with SAMtools using the samtools view and markdup functions, respectively. Mapped, filtered, and deduplicated reads were counted into genomic windows of 1Mb using the intersect function from BEDtools^59^. CNV tracks were visualized using the IGV browser^60^.

For standard Repli-seq data, sequencing reads were processed as for CNV (trimmed for adapter sequences, aligned with Burrows-Wheeler Aligner, and PCR duplicates removed). Mapped, filtered, and deduplicated reads were counted into genomic windows of distinct size (from 5 to 100 kb) using the intersect function from BEDtools^59^. Downstream processing, including quantile normalization and LOESS smoothing, and data visualization, was performed in R^61^ as previously described^22,25^ and IGV browser^60^.

For PARTAGE Repli-seq data, binned rpkm data for each sample was obtained as above and further processed into 50 kb bins. For each set of samples the log2((sample/G1_sample)+1) was calculated. Data was scaled linearly from 0 to 100 for visualization. Bedgraphs were generated and displayed using the IGV browser^60^.

For collapsed Repli-seq from PARTAGE data, fastq files were concatenated for S1, S2 as the early sample and S3 and S4 as the late sample. Data was then processed as above, including loess smoothing and quantile normalization for comparison to standard Repli-seq data.

For RNA-seq data, raw read counts were trimmed using Trimmomatic^62^, aligned using HISAT2^63^, and alignment filtering and counts using SAMtools. Raw counts were converted to TPM counts by calculating reads per kilobase (RPK) and scaling by the sum of RPK/1,000,000. The log_2_(TPM+1) was used for downstream analyses. For analysis of expression patterns, samples were median centered, low expressed genes (TPM<1) and all non-coding genes were removed, and batch correction was calculated using ComBat^45^. Mean values were calculated across samples for each timepoint. Only genes expressed in at least two samples and with a fourfold TPM change were considered. K-means clustering was performed on the remaining genes with a cluster count of two. Gene set enrichment was performed using gseapy^64^ and Enrichr^65^ on the following terms: ’ChEA_2022’, ’Chromosome_Location’, ’ENCODE_and_ChEA_Consensus_TFs_from_ChIP-X’, ’GO_Biological_Process_2025’, ’GO_Cellular_Component_2025’, ’GO_Molecular_Function_2025’, ’Reactome_Pathways_2024’. Significantly enriched terms were filtered by gene count (>1) and adjusted Benjamini-Hochberg p-value <0.05. For each category, the top 10 most significant gene categories were plotted using seaborn.

For comparisons of RNA and RT data, plots were created comparing the number of bins with expressed genes compared to their corresponding replication timing values. For each sample, genome wide RT data were sorted by value and fit into 100 bins. For each bin, gene expression values were binarized into 0s and 1s based on log2(TPM) expression > 1. A linear regression model was fit to the data using statsmodels^66^.

## DATA AVAILABILITY

### Lead contact

Requests for further information, resources, and reagents should be directed to and will be fulfilled by the lead contact, Juan Carlos Rivera-Mulia.

### Materials availability

This study did not generate new unique reagents.

### Data avvailability

All raw sequencing data has been deposited in the NCBI Sequence Read Archive (SRA). Processed data is available at the Gene Expression Omnibus at (http://www.ncbi.nlm.nih.gov/geo/),

## ACKNOWLEDGMENTS

This work was supported by NIH/NIGMS grants R35GM137950, R35GM137950-02S1, and R35GM137950-04S1 to J.C.R.M.; and by Regenerative Medicine Minnesota RMM-091621-DS-006 to J.C.R.M.

## AUTHOR CONTRIBUTIONS

Conceptualization: J.C.R.M. Methodology: J.C.R.M. Investigation: L.S.S.M., Q.D., S.M.N., and J.C.R.M. Formal analysis: Q.D., and J.C.R.M. Visualization: Q.D., and J.C.R.M. Data curation: Q.D., and J.C.R.M. Writing - original draft: L.S.S.M., Q.D., and J.C.R.M. Writing - review & editing: L.S.S.M., Q.D.,and J.C.R.M. Funding Acquisition: J.C.R.M. Resources: J.C.R.M. Supervision: J.C.R.M. Project Administration: J.C.R.M.

