## Supplementary Information for "PARTAGE: Parallel analysis of replication timing and gene expression"

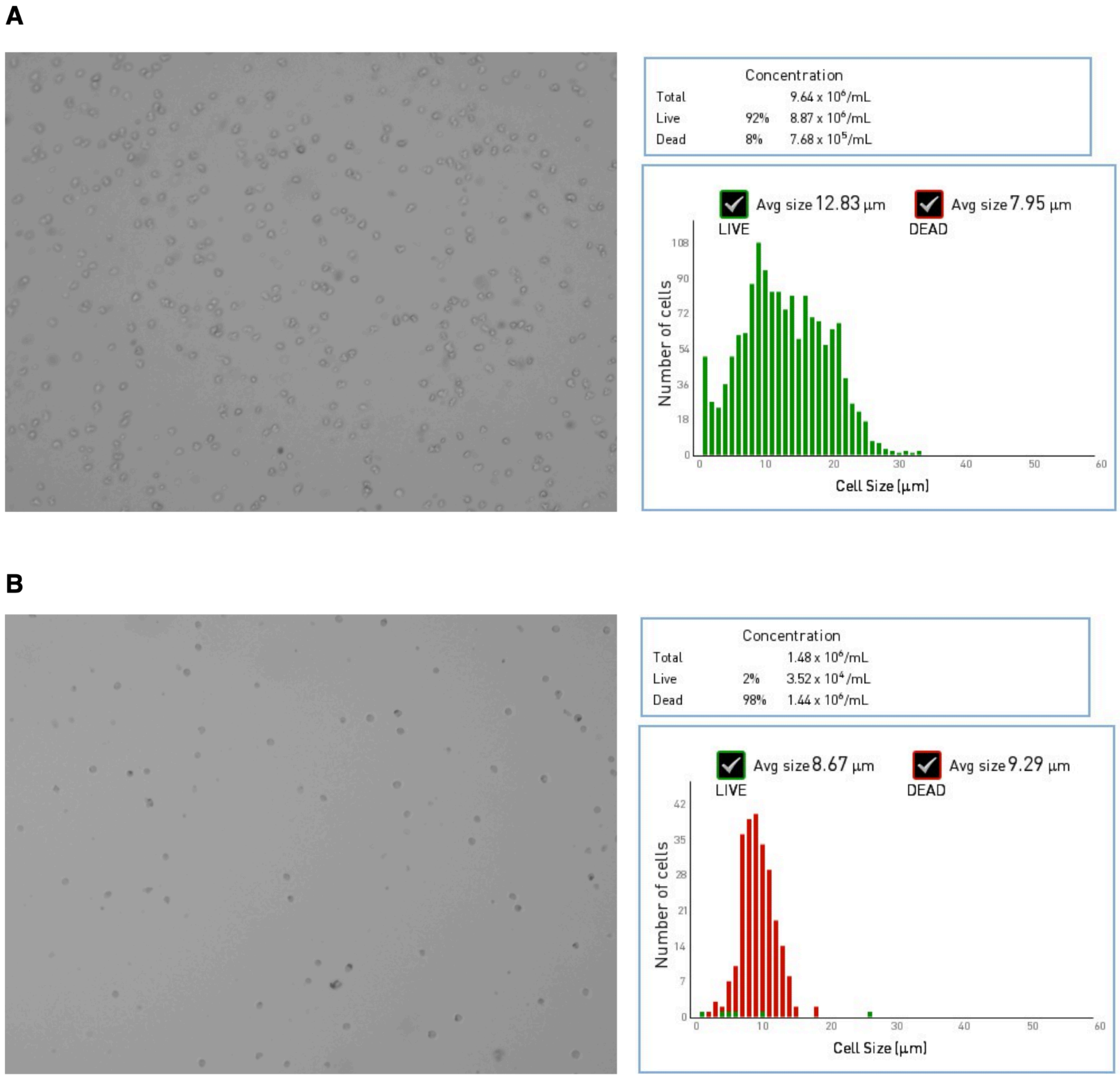


**Supplementary Figure 1**. A) Viability analysis after dissociation of HepG2 cells. B) Intact nuclei and lysis efficiency estimation. Cell viability assays were performed based on trypan blue staining on an automated cell counter (Invitrogen CountessII-FL).

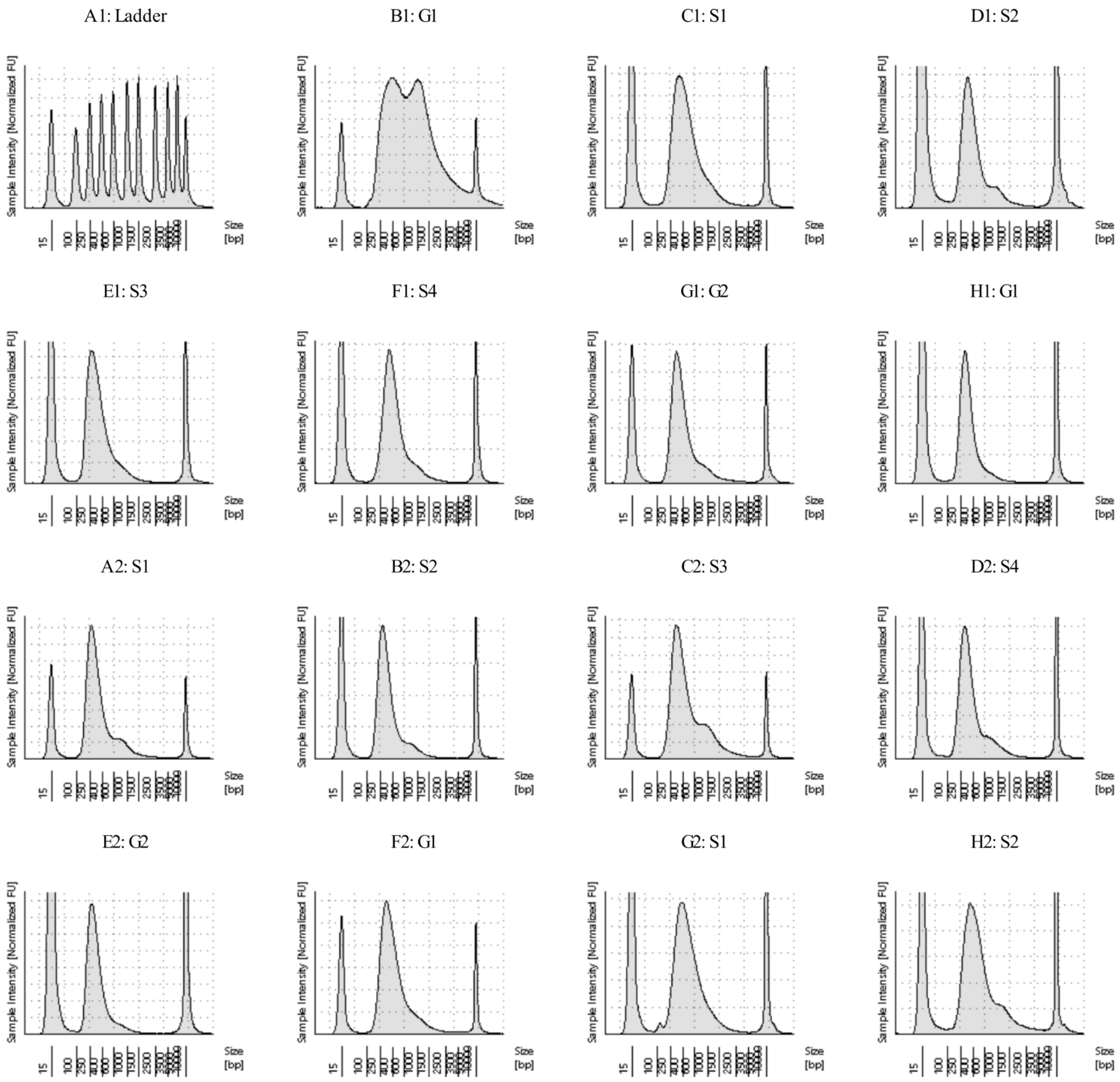


**Supplementary Figure 2**. Size distribution analysis of PARTAGE Repli-seq libraries performed on TapeStation with a High Sensitivity DNA ScreenTape.

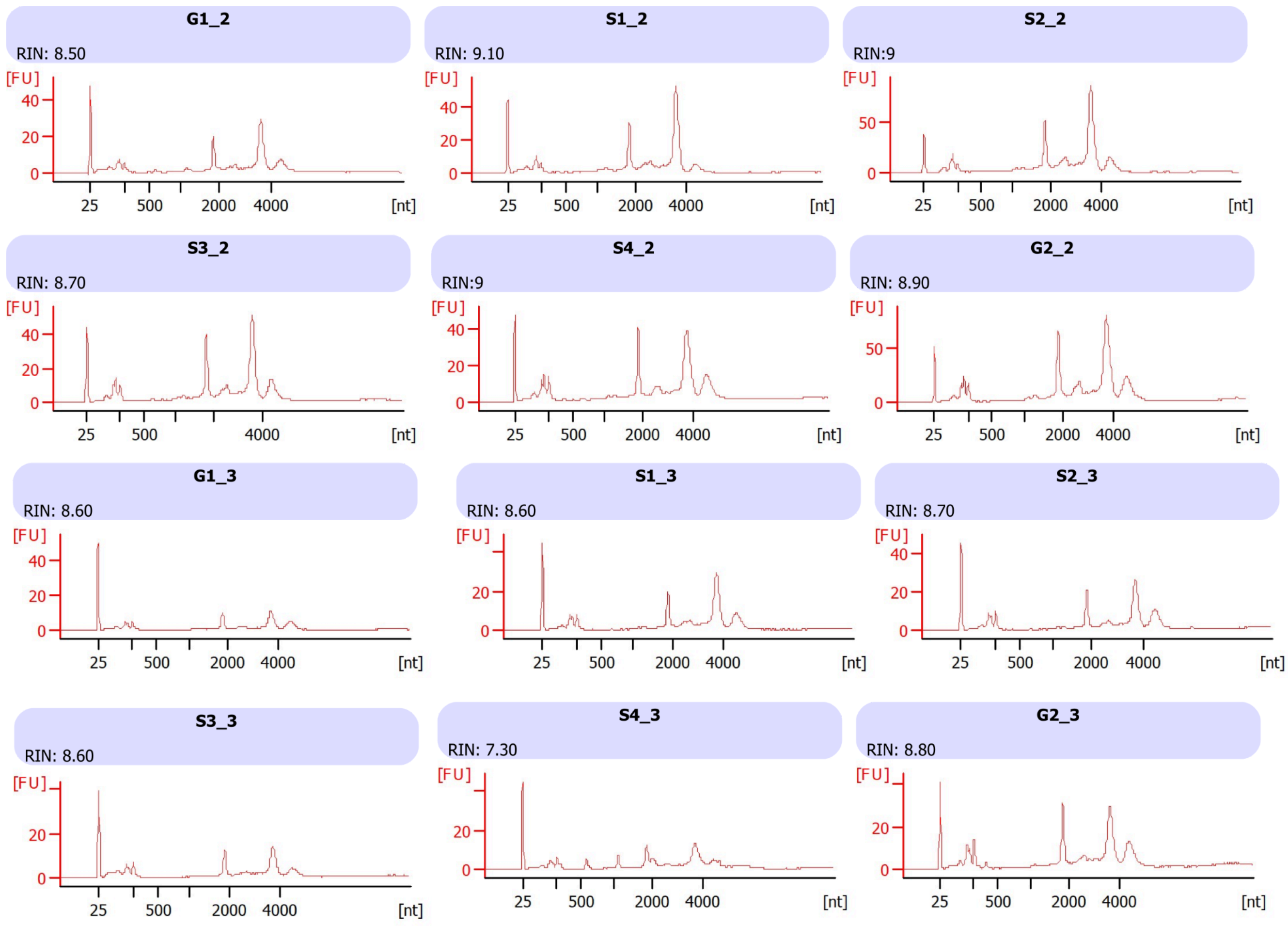


**Supplementary Figure 3**. Size distribution analysis of PARTAGE RNA-seq libraries performed on Bionalyzer with RNA 6000 Nano Kit.

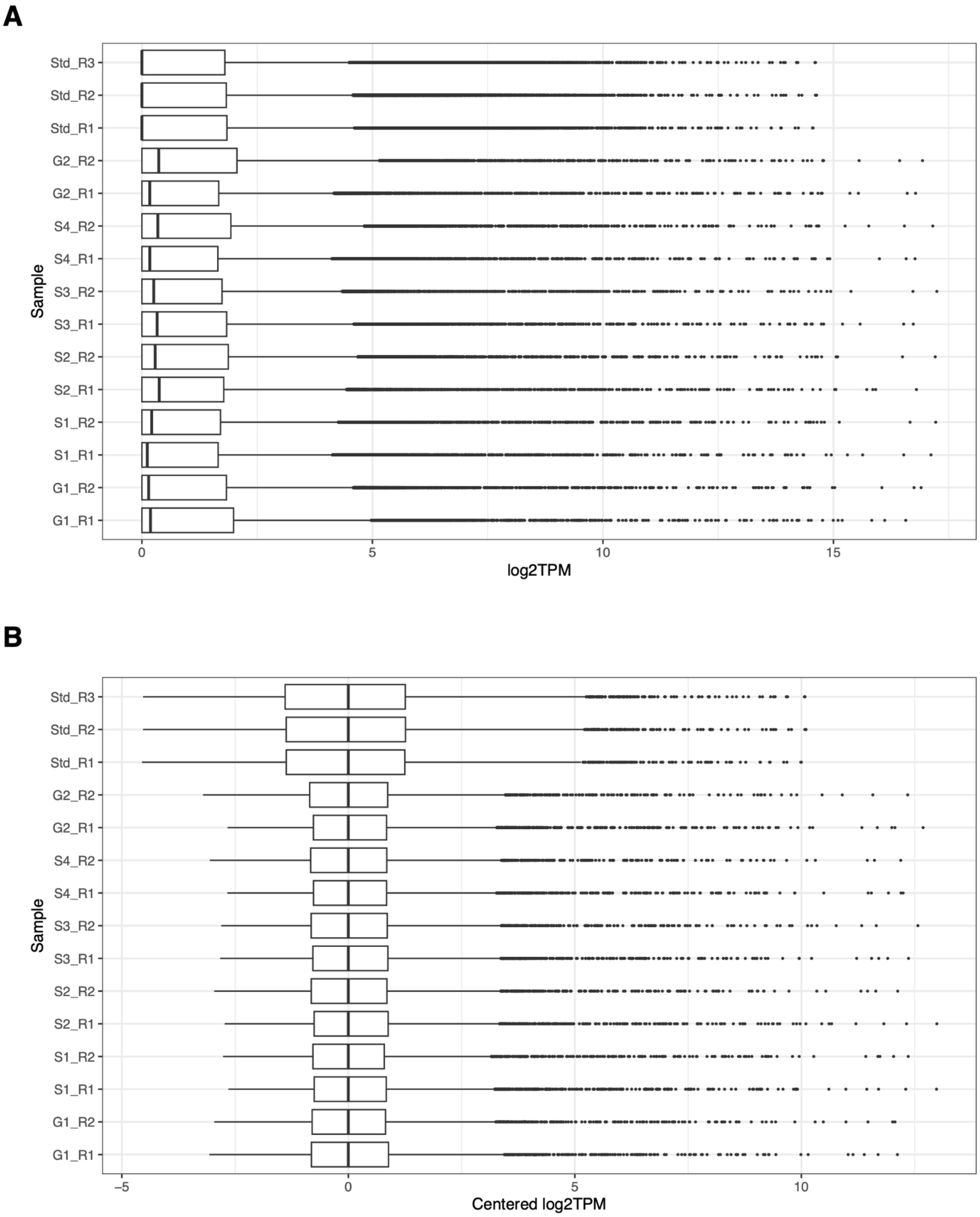


**Supplementary Figure 4**. Per sample gene expression distributions. A) Log2TPM values. B) Log2TPM values after removing non expressed genes (threshold ≥1 Log2TPM), median-centering, and batch correction using ComBat algorithm.

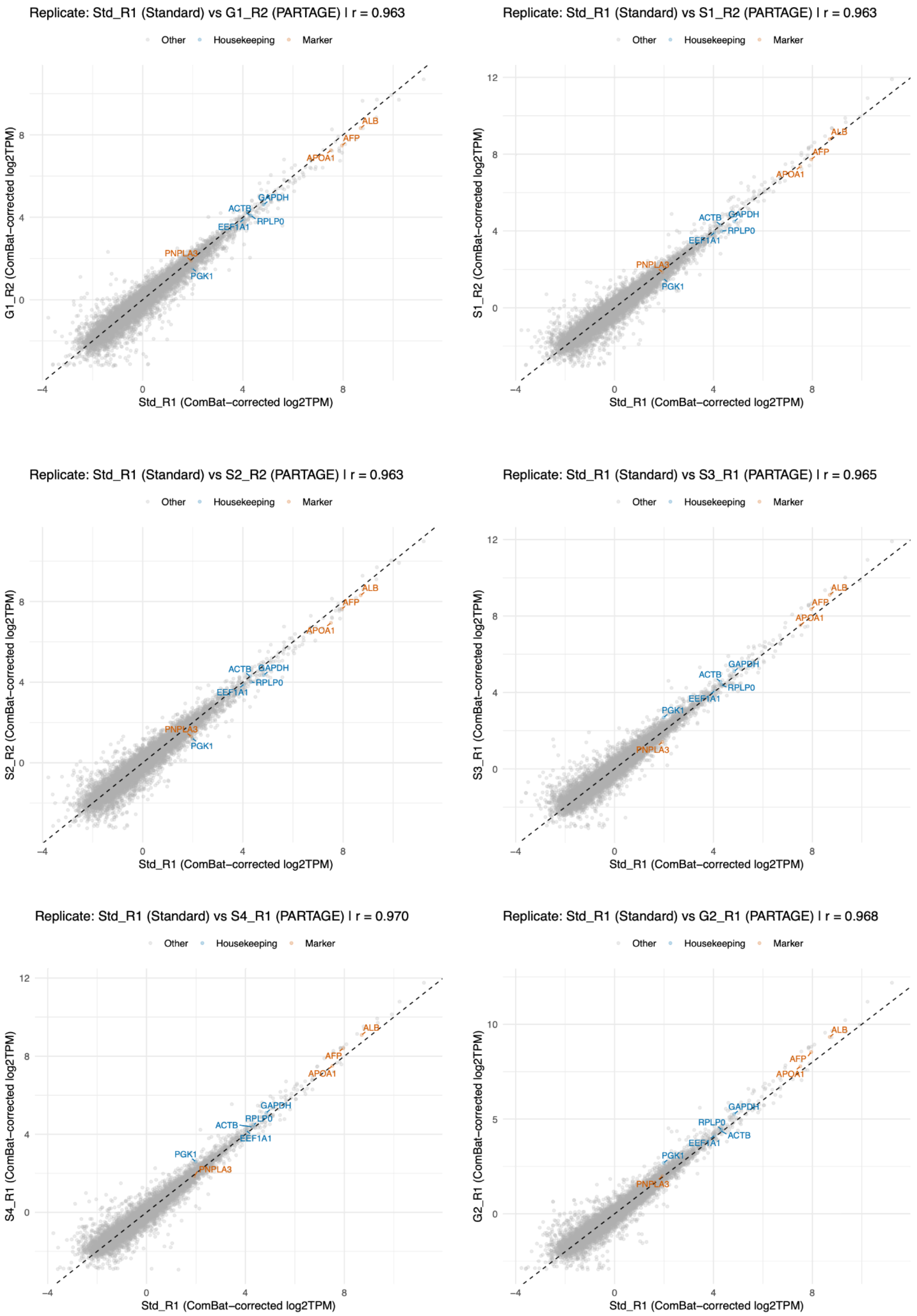


**Supplementary Figure 5**. Gene-by-gene, replicate-level scatterplots on batch-corrected gene expression values (log2TPM). Standard RNA-seq (x-axis) versus PARTAGE RNA-seq (y-axis) replicates are shown. Housekeeping genes (blue) and hepatic markers (red) are highlighted; all other genes are gray. Only genes expressed with TPM ≥ 1 in at least 2 samples were considered (11,630 protein-coding genes passed these filters).

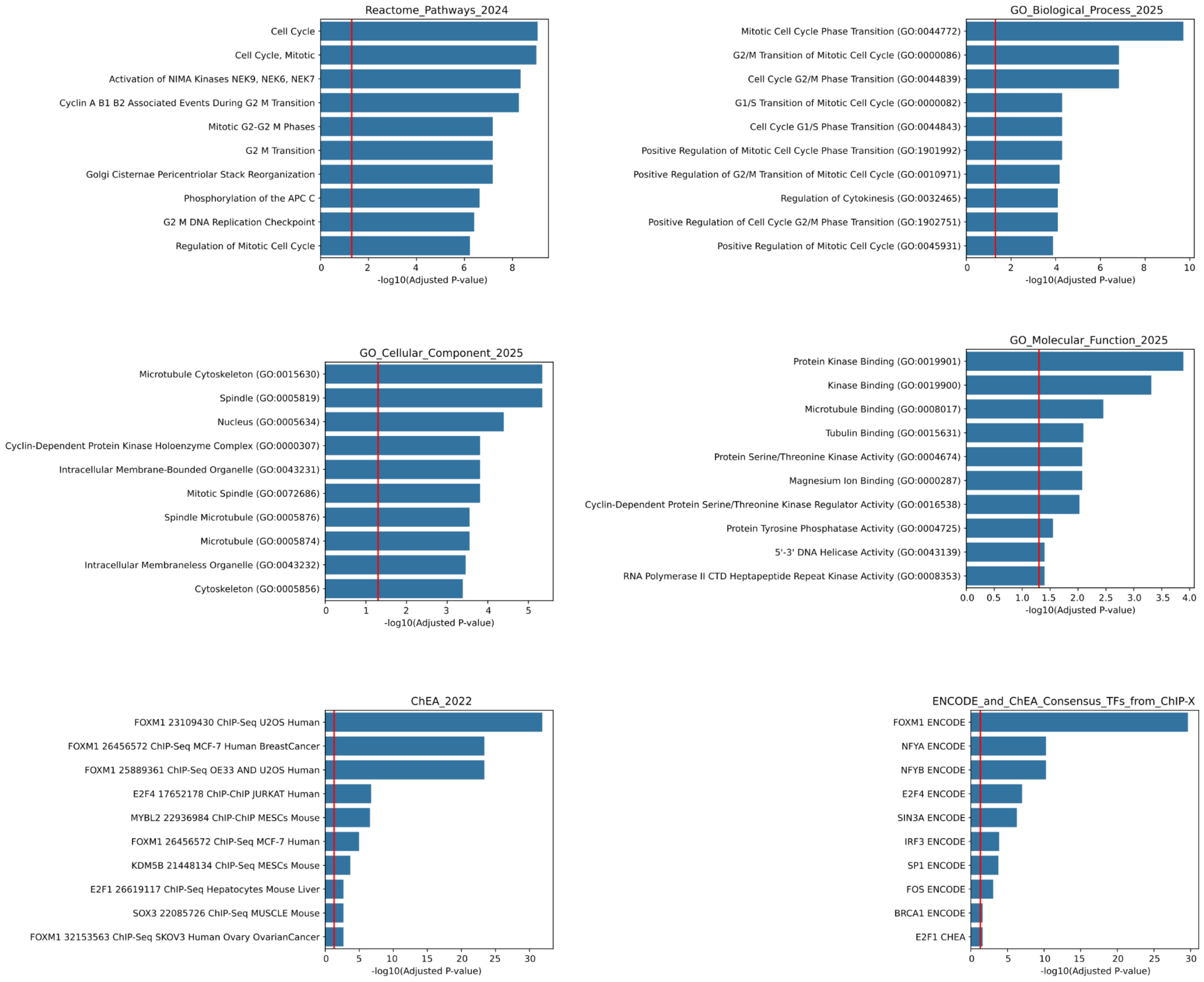


**Supplementary Figure 6**. Ontology analysis of cell cycle-regulated genes.
